# Neutral sphingomyelinase 2 regulates inflammatory responses in monocytes/macrophages induced by TNF-α

**DOI:** 10.1101/2020.05.06.080382

**Authors:** Fatema Al-Rashed, Zunair Ahmad, Reeby Thomas, Motasem Melhem, Ashley J. Snider, Lina M. Obeid, Fahd Al-Mulla, Yusuf A. Hannun, Rasheed Ahmad

## Abstract

Obesity is associated with elevated levels of TNF-α and proinflammatory CD11c monocytes /macrophages. TNF-α mediated dysregulation in the plasticity of monocytes/macrophages is concomitant with pathogenesis of several inflammatory diseases, including metabolic syndrome, but the underlying mechanisms are incompletely understood. Since neutral sphingomyelinase 2 (nSMase2; product of the sphingomyelin phosphodiesterase 3 gene, *SMPD3*) is a key enzyme for ceramide production involved in inflammation, we investigated whether nSMase2 contributed to the inflammatory changes in the monocytes/macrophages induced by TNF-α. In this study, we demonstrate that the disruption of nSMase activity in monocytes/macrophages either by chemical inhibitor GW4869 or small interfering RNA (siRNA) against SMPD3 results in defects in the TNF-α mediated expression of CD11c. Furthermore, blockage of nSMase in monocytes/macrophages inhibited the secretion of inflammatory mediators IL-1b and MCP-1. In contrast, inhibition of acid SMase (aSMase) activity did not attenuate CD11c expression or secretion of IL-1b and MCP-1. TNF-α-induced phosphorylation of JNK, p38 and NF-κB was also attenuated by the inhibition of nSMase2. Moreover, NF-kB/AP-1 activity was blocked by the inhibition of nSMase2. SMPD3 was elevated in PBMCs from obese individuals and positively corelated with TNF-α gene expression. These findings indicate that nSMase2 acts, at least in part, as a master switch in the TNF-α mediated inflammatory responses in monocytes/macrophages.

## INTRODUCTION

Obesity triggers low grade chronic inflammation in peripheral tissues such as adipose tissue and liver, which promotes various metabolic dysfunctions and thus contributes to the development of insulin resistance. In obese adipose tissue, there is an influx of immune cells, particularly monocytes (precursor cells of macrophages) and lymphocytes, which secrete inflammatory cytokines that promote inflammation (1). Monocytes isolated from obese humans display an inflammatory phenotype associated with the higher expression of surface marker of inflammation CD11c and higher secretion of pro-inflammatory cytokines/chemokines such as TNF-α, IL-6, IL-1b, and MCP-1 compared to monocytes from the lean individuals (2, 3). Monocytes/Macrophages are predominantly involved in immune modulation in obesity and respond to environmental signals with notable plasticity and adopt different forms and functions that can be characterized as pro-inflammatory M1 and anti-inflammatory M2. Obesity is associated with adipose tissue inflammation via increasing M1 macrophage polarization (4).

Obesity-driven changes of the adipose tissue monocytes/macrophages alter the secretion of the proinflammatory cytokines which contribute to the progression of inflammation and development of insulin resistance. Tumor necrosis factor-α (TNF-α) is a cytokine identified as a key regulator of inflammation and insulin resistance, and it is overexpressed in obese humans and rodents (5). In settings of obesity, TNF-α has been considered mainly pro-inflammatory as it activates immune cells particularly monocytes and macrophages into inflammatory state, M1 vs M2 (6). However, the mechanistic pathways by which TNF-α induces this pro-inflammatory shift in monocytes/macrophages remains elusive. Ceramide, a bioactive sphingolipid, has been implicated in the pathogenesis of obesity and insulin resistance (7). Ceramide is produced through various pathways; however, the hydrolysis of membrane sphingomyelin by sphingomyelinases (SMases) plays a predominant role in the production of ceramide. Neutral sphingomyelinase 2 (nSMase2), the product of the sphingomyelin phosphodiesterase 3 (*SMPD3*) gene, is a crucial enzyme involved in multiple cell regulatory pathways such as cell cycle arrest and exosome formation (8). TNF-α activates nSMase2 resulting in hydrolysis of sphingomyelin to ceramide (9). Here, we report that inhibition of nSMase2 by either chemical inhibition or siRNA reduced the TNF-α-mediated expression of inflammatory marker (CD11c) on monocytes/macrophages. Furthermore, inhibition of nSMase2 attenuated the TNF-α-mediated secretion of inflammatory markers IL-1b and MCP-1. The phosphorylation of JNK, p38 and NF-κB major downstream signaling molecules of TNF-α was decreased by inhibition of nSMase2. Moreover, our data show that *SMPD3* was elevated in the PBMCs of obese individuals and positively corelated with TNF-α. Collectively, our data reveal an interesting and novel role of nSMase2 in TNF-α-driven inflammation.

## MATERIALS AND METHODS

### Cell Culture

Monocytes, human monocytic THP-1 cells were purchased from American Type Culture Collection (ATCC) and grown in RPMI-1640 culture medium (Gibco, Life Technologies, Grand Island, USA) supplemented with 10% fetal bovine serum (Gibco, Life Technologies, Grand Island, NY, USA), 2 mM glutamine (Gibco, Invitrogen, Grand Island, NY, USA), 1 mM sodium pyruvate, 10 mM HEPES, 100 ug/ml Normocin, 50 U/ ml penicillin and 50 μg/ml streptomycin (P/S; (Gibco, Invitrogen, Grand Island, NY, USA). They were incubated at 37°C (with humidity) in 5% CO_2_. NF-kB reporter monocytic cells (THP-1-XBlue) stably expressing a secreted embryonic alkaline phosphatase (SEAP) reporter inducible by NF-κB were purchased from InvivoGen (InvivoGen, San Diego, CA, USA). -. THP-1-XBlue cells were cultured in complete RPMI medium with the addition of zeocin (200 μg/ml) (InvivoGen, San Diego, CA, USA). Prior to stimulation, monocytes were transferred to normal medium and plated in 12-well plates (Costar, Corning Incorporated, Corning, NY, USA) at 1 × 10^6^ cells/well cell density unless indicated otherwise.

#### PBMCs collection and monocyte purification

Human peripheral blood (40 ml) samples were collected from study participants in EDTA vacutainer tubes. All participants had provided written informed consent, and the study was conducted in accordance with the ethical principles of the Declaration of Helsinki and approved (04/07/2010; RA-2010-003) by the Ethical Review Committee of Dasman Diabetes Institute, Kuwait. Physical characteristics of the study participants are shown in s Table 1. PBMCs were isolated by using Histo-Paque density gradient method as we described earlier (10). PBMCs were plated in 6-well plates (Costar, Corning Incorporated, Corning, NY, USA) at 3 × 10^6^ cells/well in starvation medium for 3 hours at 37°C. Non-adhered cells were removed, and monocytes that had adhered to the plate were washed with culture media without serum and incubated for 24 hrs in RPMI with 2% Fetal Bovine serum.

**Table 1.**
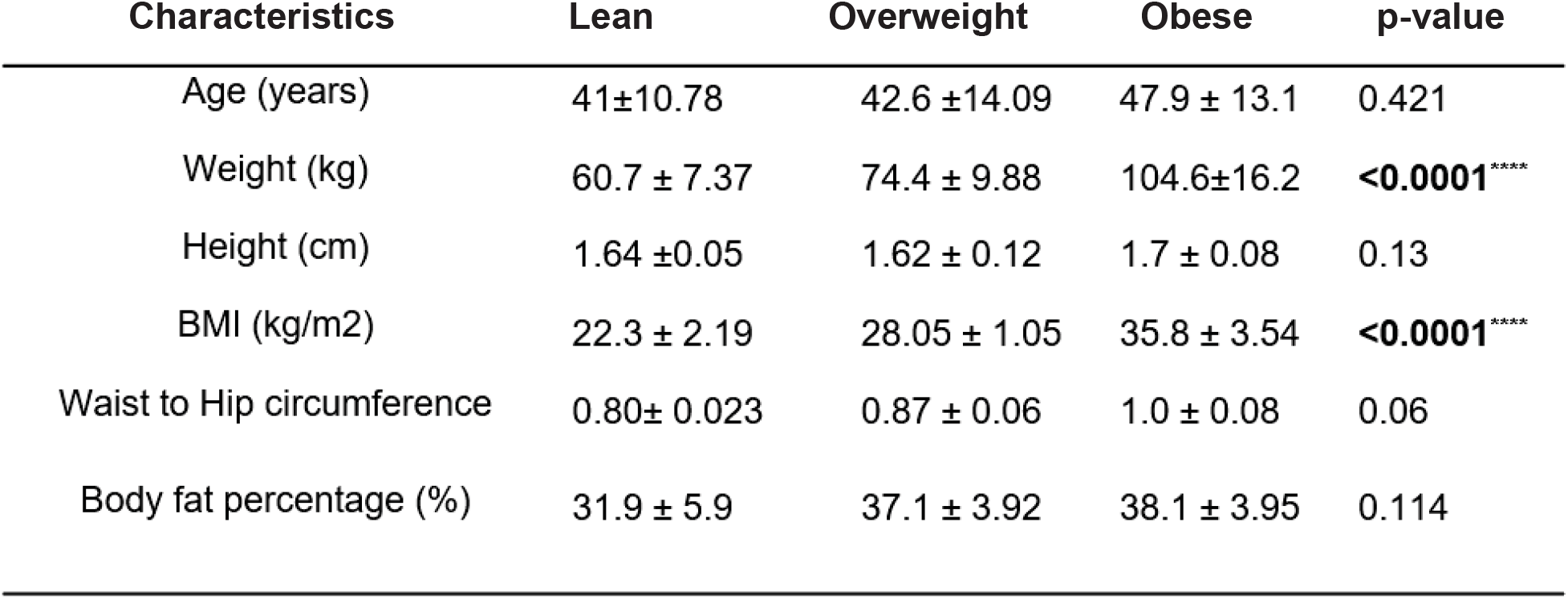
Physical characteristics of the study participants

### Cell stimulation

Monocytes were plated in 12-well plates (Costar, Corning Incorporated, Corning, NY, USA) at 1 × 10^6^ cells/well concentration unless indicated otherwise. Cells were pre-treated with Neutral sphingomyelinase nSMase2 specific inhibitor (GW4869) (Sigma D1692, 10uM/ml), then stimulated with TNF-α (10ng/ml) or Vehicle (0.1% BSA) for 2 hours at 37°C. Cells were harvested for RNA isolation. For measuring secretion of IL-1b and MCP-1 into media, TNF-α stimulation was carried out for 12 hours, and culture media were collected for analysis of IL-1b and MCP-1. For MAPK’s and NF-kB signaling pathway analysis, cultures were treated with the inhibitor as stated above then stimulated with TNF-α or BSA vehicle for 10-15 min.

### Real time quantitative RT-PCR

Total RNA was extracted using RNeasy Mini Kit (Qiagen, Valencia. CA, USA) per the manufacturer’s instructions. The cDNA was synthesized using 1μg of total RNA using high capacity cDNA reverse transcription kit (Applied Biosystems, Foster city, CA, USA). Real-time PCR was performed on 7500 Fast Real-Time PCR System (Applied Biosystems, Foster City, CA, USA) using TaqMan® Gene Expression Master Mix (Applied Biosystems, Foster city). Each reaction contained 50ng cDNA that was amplified with Inventoried TaqMan Gene Expression Assay products (SMPD3: Assay ID:Hs00920354_m1; SMPD2:Assay ID: Hs00906924_g1; SMPD1:Assay ID: Hs03679346_g1; CD11c: Assay ID: Hs00174217_m1; GAPDH: Hs03929097_g1;). The threshold cycle (Ct) values were normalized to the house-keeping gene GAPDH, and the amounts of target mRNA relative to control were calculated with ΔΔCt-method(11, 12). Relative mRNA expression was expressed as fold expression over average of control gene expression. The expression level in control treatment was set at 1. Values are presented as mean ± SEM. Results were analyzed statistically; P< 0.05 was considered significant.

### Extracellular staining -flow cytometry

Monocytic cells were seeded in 24 well plate at 0.5×10^5^ cell/ml in serum free media overnight. Cells were treated with nSMase inhibitor GW4869 (10uM/ml) for one hour or 0.01% DMSO (vehicle) then subjected to stimulation with TNF-α (10ng/ml) or BSA (vehicle) for 6 hours. Monocytic cells (1×10^6^ cells) were resuspended in FACS staining buffer (BD Biosciences) and blocked with human IgG (Sigma; 20μg) for 30 minutes on ice. Cells were washed and resuspended in 100 ul of FACS buffer and incubated with CD11c (S_HCL-3)-PE (cat# 347637; BD Biosciences) or CD11c PE-Cy7 (cat # 117317; BD Biosciences) on ice for 30 minutes. Cells were washed three times with FACS buffer and resuspended in 2% paraformaldehyde. Cells were centrifuged and resuspended in FACS buffer for FACS analysis (FACSCanto II; BD Bioscience, San Jose, USA). FACS data analysis was performed using BD FACSDiva™ Software 8 (BD Biosciences, San Jose, USA).

### Intracellular Staining -flow Cytometry

Flow cytometry analysis was used to investigate the expression of signaling pathway markers. Briefly, cells were seeded in 24 well plate at 0.5×10^5^ cells/ml in serum free media overnight. Cells were treated with nSMase inhibitor GW4869 or DMSO (vehicle) then subjected to stimulation with TNF-α (10ng/ml) or BSA (vehicle) for 10 min. After stimulation, cells were collected and washed. Cells were then incubated with fixation/permeabilization buffer (cat# 00-5523-00, eBioscience, San Diego, CA, USA) for 20 min in 4°C, followed by washing and staining with mouse anti-human p-NF-kB P65-PE (cat # 558423; BD Biosciences) and Alexa Fluor®647 mouse anti-IκBα (cat # 560817; BD Biosciences) for 30 min. The cells were then washed and resuspended in PBS supplemented with 2% FCS for FACS analysis (FACSCanto II; BD Bioscience, San Jose, USA). FACS data analysis was performed using BD FACSDivaTM Software 8 (BD Biosciences, San Jose, USA).

### IL-1b and MCP-1 determination

Secreted IL-1b and MCP-1 proteins in supernatants of monocytic cells stimulated with TNF-α were quantified using sandwich ELISA following the manufacturer’s instructions (R&D systems, Minneapolis, USA).

### Small interfering RNA (siRNA) transfections

Monocytes were washed and resuspended in 100 ul of nucleofector solution provided with the Amaxa Noclecfector Kit V and transfected separately with siRNA-n-SMase (Santa Cruz Biotechnology, INK. SC-106277), scramble (control) siRNA (30nM; OriGene Technologies, Inc. MD, USA, USA), and pmaxGFP (0.5 ug; Amaxa Noclecfector Kit V for THP-1, Lonza). All transfection experiments were performed with Amaxa Cell Line Nucleofector Kit V for monocytic cells(Lonza, Germany) by using Amaxa Electroporation System (Amaxa Inc, Germany) according to the manufacturer’s protocol (13). After 36 hours of transfection, cells were treated with TNF-α for 2 hours. For knock down of nSMase2 in human primary monocyte cultures, cells were transfected with 20 nM of the siRNA using Viromer Blue (lipocalyx, Halle, Germany) according to the manufacturer’s instructions. Cells were harvested for RNA isolation and staining study. nSMase gene knock down level was assessed by Real Time-PCR using SMPD3 gene-specific primer probes.

### Measurement of NF-κB activity

NF-kB reporter monocytes (THP-1 XBlue; InvivoGen, San Diego, CA) are stably transfected with a reporter construct expressing a secreted embryonic alkaline phosphatase (SEAP) gene under the control of a promoter inducible by the transcription factors NF-κB. Upon stimulation, NF-κB is activated and subsequently the secretion of SEAP is stimulated. Cells were stimulated with TNF-α (10 ng/ml) for 6-12 hours at 37°C. Levels of SEAP were detected in the culture media after 3 hours incubation of supernatants with Quanti-Blue solution (InvivoGen, San Diego, CA, USA) at 650nm wavelength by ELISA reader.

### Statistical Analysis

Statistical analysis was performed using GraphPad Prism software (La Jolla, CA, USA). Data are shown as mean ± standard error of the mean, unless otherwise indicated. Unpaired Student t-test and one-way ANOVA followed by Tukey’s test were used to compare means between groups. For all analyses, data from a minimum of three sample sets were used for statistical calculation. P value <0.05 was considered significant. Ns: no significance, *P < 0.05, ** P<0.01, *** P< 0.001 and **** P < 0.0001)

## RESULTS

### Inhibition of the nSMase2 attenuates TNF-α induced inflammatory responses in monocytic cells

nSMase2 is involved in many pathophysiological inflammatory processes (14). However, its role in the TNF-α-mediated plasticity of monocytes/macrophages is not defined yet. In the settings of obesity, CD11c is abundantly expressed in monocytes and macrophages (2). Therefore, we questioned whether nSMase2 was required for TNF-α-mediated activation of primary monocytes, using GW4869 to inhibit nSMase activity. Pretreatment with the nSMase2 inhibitor GW4869, followed by exposure to TNF-α, caused a significant suppression in the expression of the phenotypic inflammatory marker (CD11c) at both mRNA and protein levels in human primary monocytes **(Fig. 1A-C).** Daunorubicin (DNR) has been shown to activate nSMase2 (15); therefore, we asked whether DNR induced changes in the expression of inflammatory markers in monocytic cells. Our data show that DNR induces inflammatory change in monocytes that was comparable to TNF-α, manifested by increased CD11c expression (**Fig. 1A-C**).

**Figure 1.**
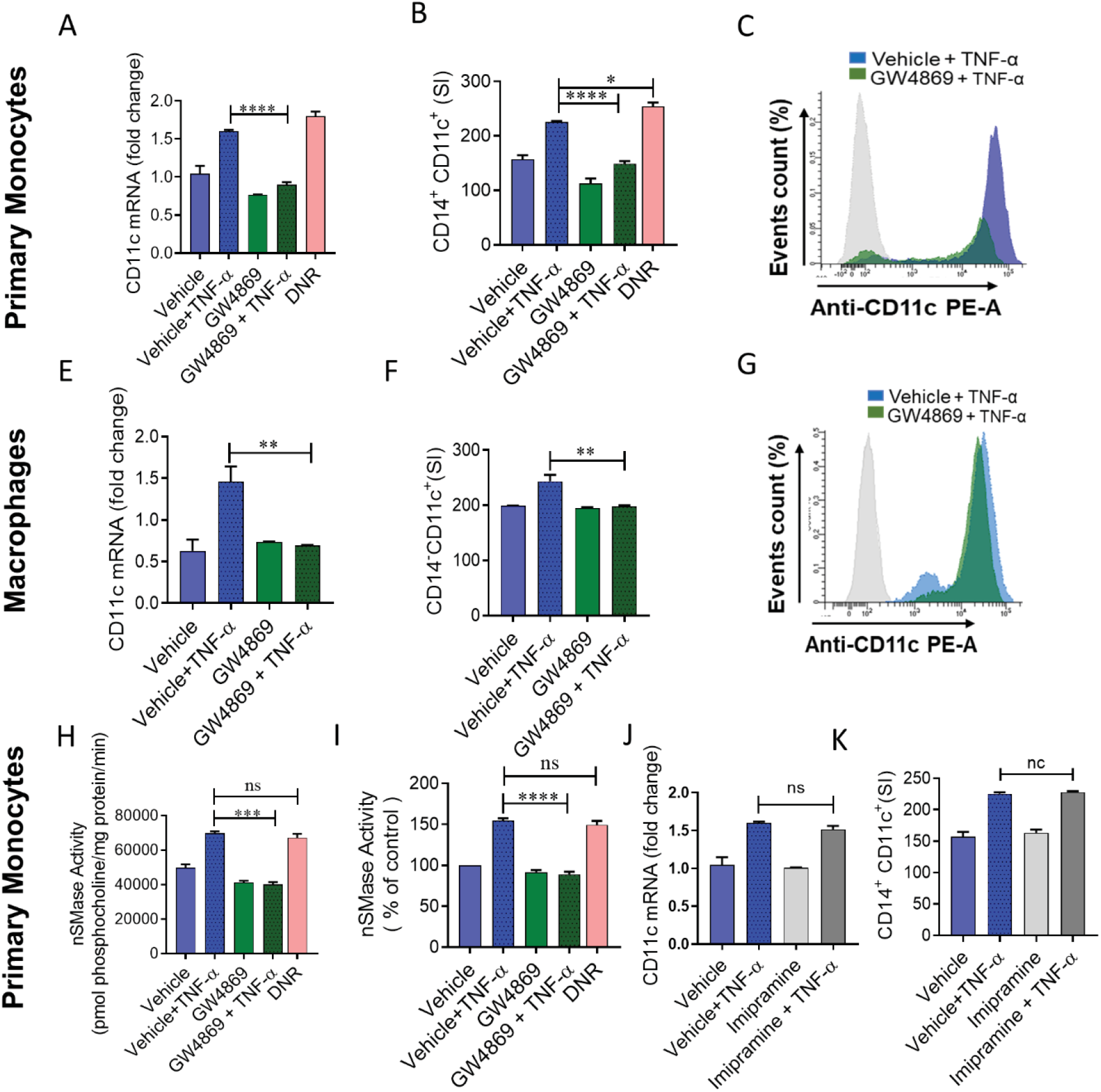
nSMase inhibition blocks TNF-α mediated pro-inflammatory changes in human monocytes/macrophages. Primary monocytes were pretreated with nSMase inhibitors (GW4869: 10uM), nSMase agonist (DNR; 80nM) or vehicle for 1 hour and then incubated with TNF-α for 2 hours. Cells were harvested, and mRNA of CD11c was determined by real time RT-PCR **(A).** After 6 hours treatment with TNF-α, cells were stained with antibodies against CD14 and CD11c along with matched isotype controls. Surface expression of CD14^+^CD11c^+^ was assessed by flow cytometry, **(B)** data are presented as a bar graph of mean staining index, and **(C)** representative histogram. Macrophages derived from monocytes were pre-incubated with (GW4869: 10uM) for 1 hour and then treated with TNF-α. **(E)** CD11c mRNA expression and **(F and G)** surface expression of CD14^+^CD11c^+^ were assessed by flow cytometry. Monocytic cells (Primary monocytes) were pretreated with nSMase inhibitor (GW4869: 10uM) or vehicle for 1 hour and then incubated with TNF-α for 10min. **(H)** nSMase activity measured in in equal amount of protein (20ug); (I) nSMase activity as percentage of expression to vehicle. Primary monocytes were pretreated with aSMase inhibitor (lim: 10uM) or vehicle for 1 hour and then incubated with TNF-α for 2 hours. **(J and K)** Cells were harvested, and mRNA of CD11c and surface expression of CD14^+^CD11c^+^ cells were determined. All data are expressed as mean ± SEM (n ≥ 3). *p≤0.05, **p≤0.01, ***p≤0.001, ****p≤0.0001 versus vehicle.

Since macrophages are key regulators of adipose tissue functions during metabolic inflammation, we further asked whether chemical inhibition of nSMase2 attenuates macrophage expression of CD11c. Using macrophages derived from primary monocytes, we found that the presence of GW4869 inhibited TNF-α-induced CD11c upregulation in macrophages **(Fig.1E-G,).** A similar observation was seen in THP1 human monocytic cells **(Supplemental Fig. 1A-C).** Next, we determined nSMase activity in monocytic cells treated with TNF-α or DNR. In this regard, we found that nSMase activity was higher in cells treated with TNF-α or DNR compared to the cells treated with vehicle. Furthermore, GW4869 significantly inhibited TNF-α mediated nSMase activity in monocytic cells (**Fig. 1H and I).**

TNF-α has also been reported to upregulate aSMase activity and subsequently modulate inflammatory signaling pathways (16). Therefore, we wanted to know whether there was any involvement of aSMase in the inflammatory responses induced by TNF-α. To this end, cells were treated with imipramine (a functional inhibitor of aSMase (17) followed by treatment with TNF-α. The results indicated that there was no change after imipramine treatment in TNF-α mediated inflammatory responses in monocytic cells **(Fig. 1 J and K)**. These results rule out the possible involvement of aSMase in TNF-α mediated inflammatory responses. Collectively our results show that nSMase is involved in the TNF-α mediated upregulation of CD11c expression in human monocytic cells and macrophages.

### nSMase inhibition reduces TNF-α mediated IL-1b and MCP-1 production

Activated monocytes/macrophages contribute to the pathogenesis of several diseases by secreting inflammatory cytokines including IL-1b and MCP-1 (18). Next, we asked whether IL-1b and MCP-1 production by TNF-α activated primary monocytes or THP1 monocytic cells was reduced by the inhibition of nSMase2. The results showed that inhibition of nSMase2 significantly reduced the production of IL-1b and MCP-1 by TNF-α activated primary monocytes (**Fig. 2A and B**), macrophages (**Fig. 2 C and D**) and THP1 monocytic cells (**Fig. 2 E and F**). However, inhibition of aSMase activity by imipramine did not suppress TNF-α induced IL-1b and MCP-1(**Fig. 2A-B and 2E-F**). DNR, which activates nSMase2, showed similar production of cytokines as noted in case of TNF-α stimulation (**Fig. 2A-B and 2E-F)**, suggesting that induction of nSMase2 is sufficient to induce these changes. Together, these results indicate that TNF-α regulates the production of key inflammatory cytokines IL-1b and MCP-1 through the activation of nSMase.

**Figure 2.**
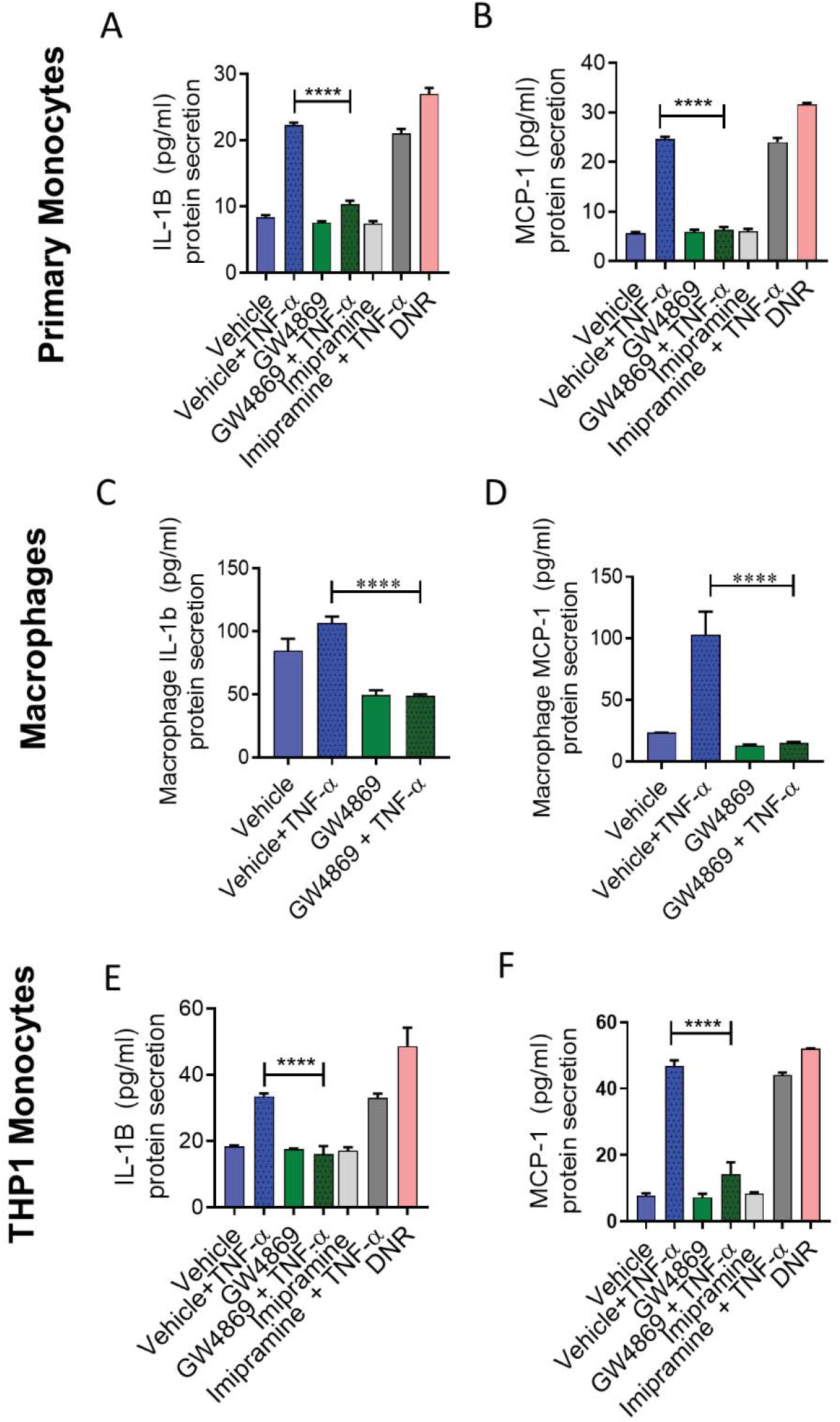
IL-1β and MCP-1 secreted by TNF-α activated monocytes are suppressed by nSMase inhibition. Monocytic cells (primary monocytes, macrophages, THP-1 cells) were pretreated with nSMase inhibitors (GW4869: 10uM), nSMase agonist (DNR: 80nM), aSMase inhibitor (Lim: 10uM) or vehicle for 1 hour and then incubated cells with absence or presence of TNF-α for 12 hours. Secreted IL-1β and MCP-1 protein in culture media was determined by ELISA. **(A and B)** IL-1β and MCP1 secreted by primary monocytes, **(C and D)** primary macrophages and **(E and F)** THP-1 cells. All data are expressed as mean ± SEM (n ≥ 3). *p≤0.05, **p≤0.01, ***p≤0.001, ****p≤0.0001 versus vehicle

### nSMase-2 deficiency impairs TNF-α mediated pro-inflammatory responses in human monocytes/macrophages

To further define the role of nSMase2 in TNF-α-induced inflammatory alteration in primary monocytes and in THP1 monocytic cells, we transfected cells with siRNA against nSMase2 (SMPD3), which achieved 50 −70% reduction in nSMase2 mRNA levels compared with scramble (control) siRNA (**Fig 3A and E**). Results showed that the expression of CD11c, at both the mRNA and protein levels, was significantly reduced in SMPD3 siRNA–transfected primary monocytes **(Fig. 3B-D**) as well as THP1 monocytes (**Fig. 3F-H**) after stimulation with TNF-α compared to the cells transfected with scramble siRNA. Next, we wanted to see whether nSMase2 deficiency in monocytic cells affected the TNF-α induced production of IL-1b and MCP-1. As expected, our results showed that monocytic cells deficient in nSMase2 failed to respond to TNF-α treatment for secretion of IL-1b and MCP-1 **(Fig. 3I-J).** Overall, the results suggest the significant involvement of nSMase2 in inflammatory responses induced by TNF-α in monocytic cells.

**Figure 3.**
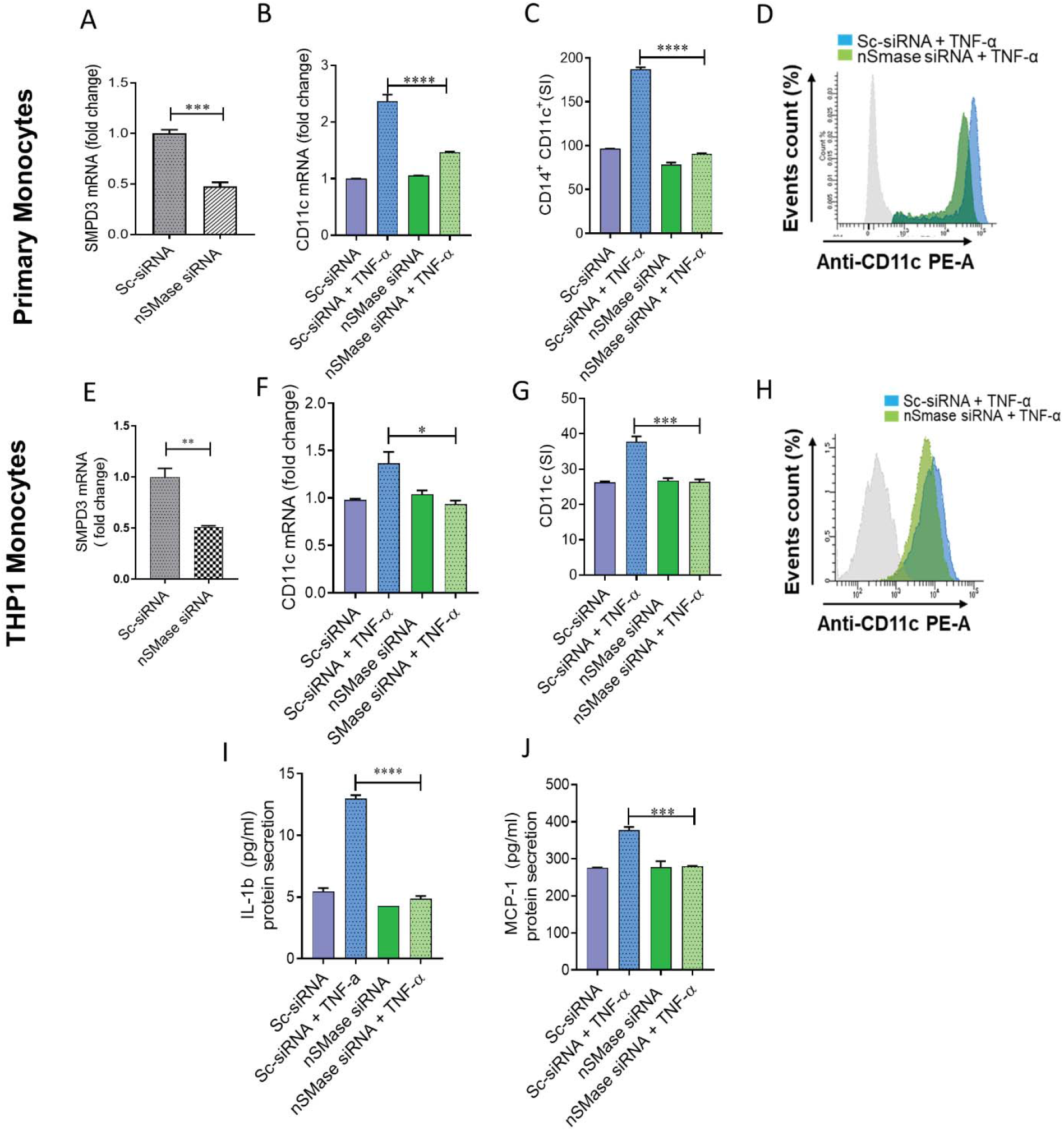
TNF-α mediated pro-inflammatory monocytic responses requires nSMase. Primary monocytes and THP-1 monocytes were transfected with scramble-siRNA (negative control; Sc) or SMPD3 siRNA and incubated for 36 hours. Real-time PCR was performed to measure **(A)** SMPD3 expression in primary monocytes and **(B)** CD11c was determined by real time RT-PCR. Cells were stained with antibodies against CD14 and CD11c along with matched isotypes and were subjected to flow cytometry analysis. **(C)** Flow cytometry data are presented as a bar graph of mean staining index of CD14^+^CD11c^+^ cells as well as **(D)** representative histogram. nSMase deficient THP-1 cells were treated with TNF-α and vehicle. **(E)** SMPD3 expression in transfected THP-1 monocytic cells. **(F)** CD11c mRNA was determined by real time RT-PCR. Cells were stained with antibody against CD11c along with matched isotype controls and assessed by flow cytometry. **(G)** Flow cytometry data are presented as a bar graph of mean staining index as well as **(H)** representative histogram. (I-J) Secreted IL-1b and MCP-1 by nSMase2 deficient cells. All data are expressed as mean ± SEM (n ≥ 3). *p≤0.05, **p≤0.01, ***p≤0.001, ****p≤0.0001 versus vehicle.

### MAPKs and NF-kB activation signal downstream of nSMase2 in TNF-α mediated inflammation

MAPK and NF-kB signaling pathways function downstream of TNFR activation. Therefore, we questioned whether nSMase is involved in the TNF-α mediated activation of MAPK and NF-κB signaling pathways. To gain insight into the nSMase function in the TNF-α-induced activation of the MAPK and NF-kB signaling pathways, we treated cells with the nSMase inhibitor (GW4869) prior to TNF-α treatment. The results showed that inhibition of nSMase significantly reduced the TNF-α mediated phosphorylation of ERK1/2 (**Fig. 4A**), p38 (**Fig. 4B**), JNK (**Fig. 4C**), c-Jun (**Fig. 4D**) and the p65 subunit of NF-kB (**Fig. 4E-H).**

**Figure 4.**
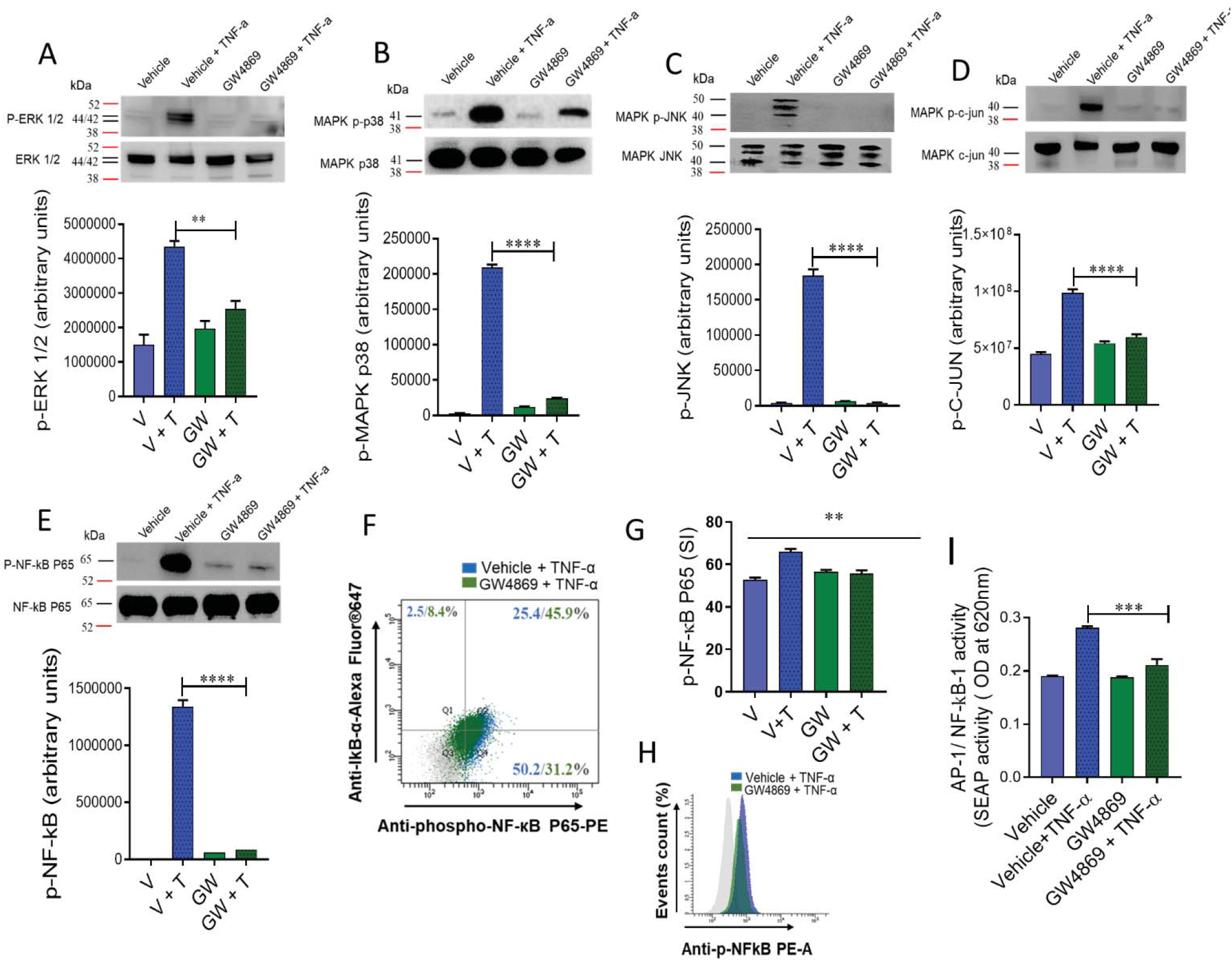
Inhibition of nSMase affects TNF-α activated MAPK and NF-κB signaling pathways in monocytic cells. Monocytic cells were pretreated with nSMase inhibitor (GW4869: 10uM) and then incubated with TNF-α for 15 minutes. Cell lysates were prepared as described in Materials and Methods. Samples were run on denaturing gels. Immune-reactive bands were developed using an Amersham ECL Plus Western Blotting Detection System (GE Healthcare, Chicago, IL, USA) and visualized by Molecular lmager® ChemiDoc™ MP Imaging Systems (Bio-Rad Laboratories, Hercules, CA, USA). **(A)** Phosphorylated proteins of ERK1/2 (p42/44), **(B)** p38 MAPK, **(C)** JNK, **(D)** c-Jun and **(E)** NF-κB are depicted in the upper panels and total respective proteins are shown in the lower panels. Phosphorylation intensity of p38 MAPK, ERK1/2, and NF-κB was quantified using Image Lab software (version 6.0.1, Bio-Rad, Hercules, CA, USA) and are presented in arbitrary units. All data are expressed as mean ± SEM (*n* ≥ 3). **** P <0.0001 versus TNF-α without respective inhibitor. NF-κB phosphorylation was also determined by flow cytometry. **(F)** Representative flow cytometry dot plots of p-NF-κB fluorescence versus total IκBα cells. **(G)** Flow cytometry data are presented as a bar graph of mean staining index as well as **(H)** representative histogram. Bar graphs depict mean values ± SEM of staining intensity (SI). P<0.05 was considered as statistically significant (*P≤0.05; **P≤0.01, ***P≤0.001, ****P≤0.0001). The data in all Figures are representative of three independent experiments. **(I)** NF-κB reporter monocytic cells were pretreated with nSMase inhibitor (GW4869: 10uM) or vehicle for 1 hour and then incubated with TNF-α for 12 hours. Cell culture media were assayed for SEAP reporter activity (degree of NF-κB activation).

NF-κB and AP-1 are downstream transcription factors of TNF-α activation. To further confirm the role of nSMase2 in TNF-α mediated activation of NF-κB/AP-1, we used NF-κB/AP-1 activity reporter human monocytic cells. The data showed that TNF-α induced higher NF-κB/AP-1 activity in the reporter cells and this activity was not seen in the cells treated with nSMase2 inhibitor (**Fig. 4I**). Together, these data support the role of nSMase2 in the TNF-α mediated activation of NF-kB and AP1 transcription factors.

### Association of TNF-α and nSMase2 expression in human PBMCs with various BMI

The *in vitro* data implicated the involvement of nSMase2 in the TNF-α mediated inflammatory responses in the human monocytes/macrophages. Next, we examined the role of nSMase2 and TNF-α in PBMCs of the obese individuals. To this end, we isolated RNA from the PBMCs of 40 individuals (lean, overweight, obese) and determined gene expression SMPD1(aSMase), SMPD2, SMPD3 and TNF-α (Fig. 5A-D). The data showed that nSMase2 (SMPD3) and TNF-α expression levels are elevated in obese individuals as compared to lean (**Fig. 5C-D).** Furthermore, nSMase2 was positively correlated with TNF-α (**Fig. 5E**). However, aSMase gene (SMPD1) expression was not elevated in PBMCs of the obese individuals as compared to lean individuals (**Fig. 5A**).and had no association with TNF-α (**Fig. 5F**). Likewise, the expression of nSMase1 (product of SMPD2 gene) was not elevated (**Fig. 5B**). Overall, nSMase2 and TNF-α show an elevated and associated expression in human PBMCs.

**Figure 5.**
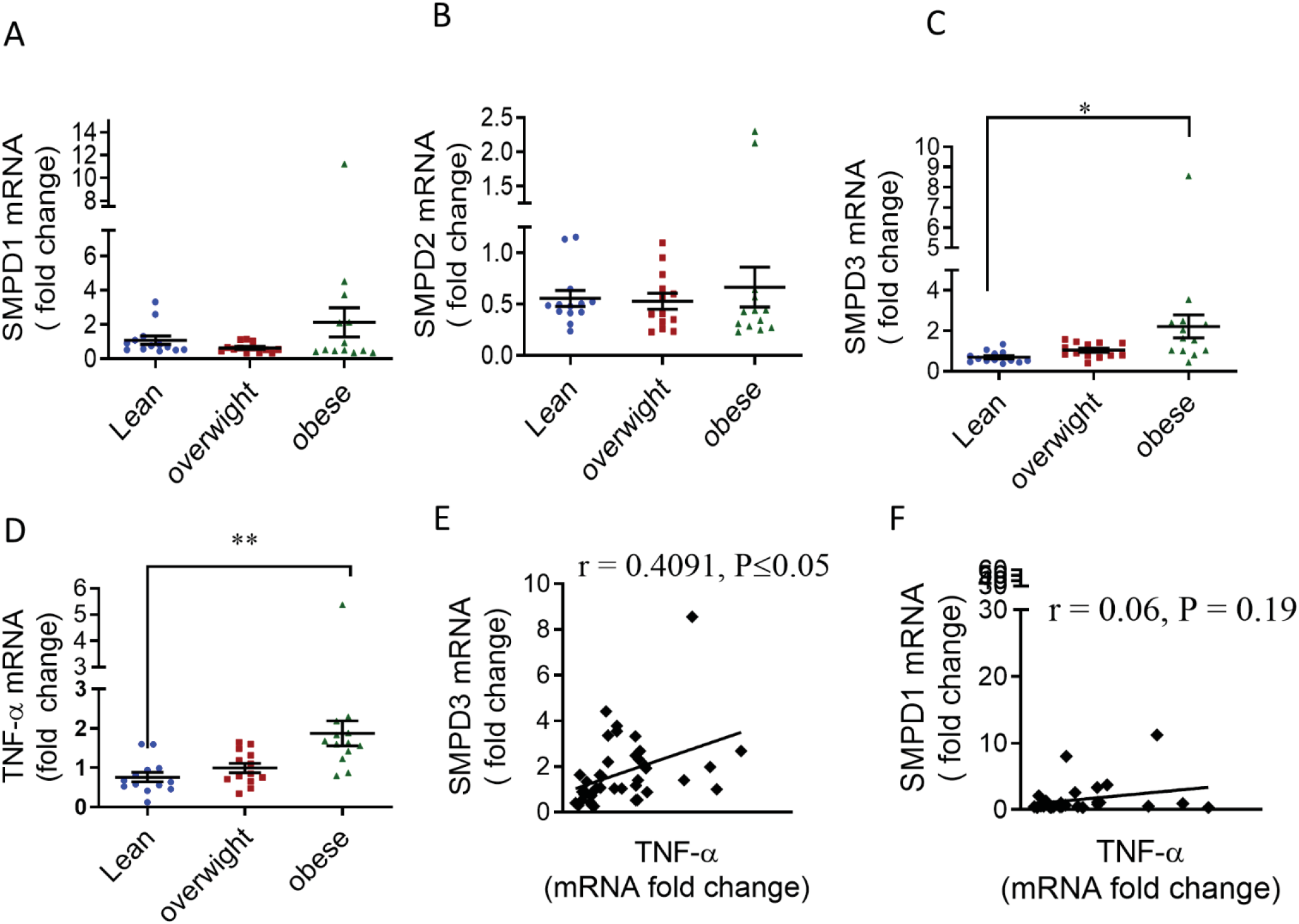
Association of elevated TNF-α with SMPD3 levels in obese human PBMCs. PBMCs were isolated from human blood samples obtained from lean (n= 13), overweight (n= 13) and obese (n=13) individuals. mRNA of *SMPD1 (aSMase), SMPD2 (nSMase), SMPD3 (nSMase)* and TNF-α were detected by real time RT-PCR and represented as fold change over controls. **(A -D)** Each dot represents the individual value of SMPD1, SMPD2, SMPD3 or TNF-α mRNA. Lines represented the mean values of PBMCs’ SMPD1, SMPD2, SMPD3 and TNF-α of each group. **(E)** Correlation of *SMPD3 and (F) SMPD1* mRNA’s with TNF-α in PBMCs of obese individuals.

To summarize the involvement of nSMase2 in TNF-α mediated inflammatory responses in human monocytic cells, a schematic illustration is presented in **Fig. 6**.

**Figure 6.**
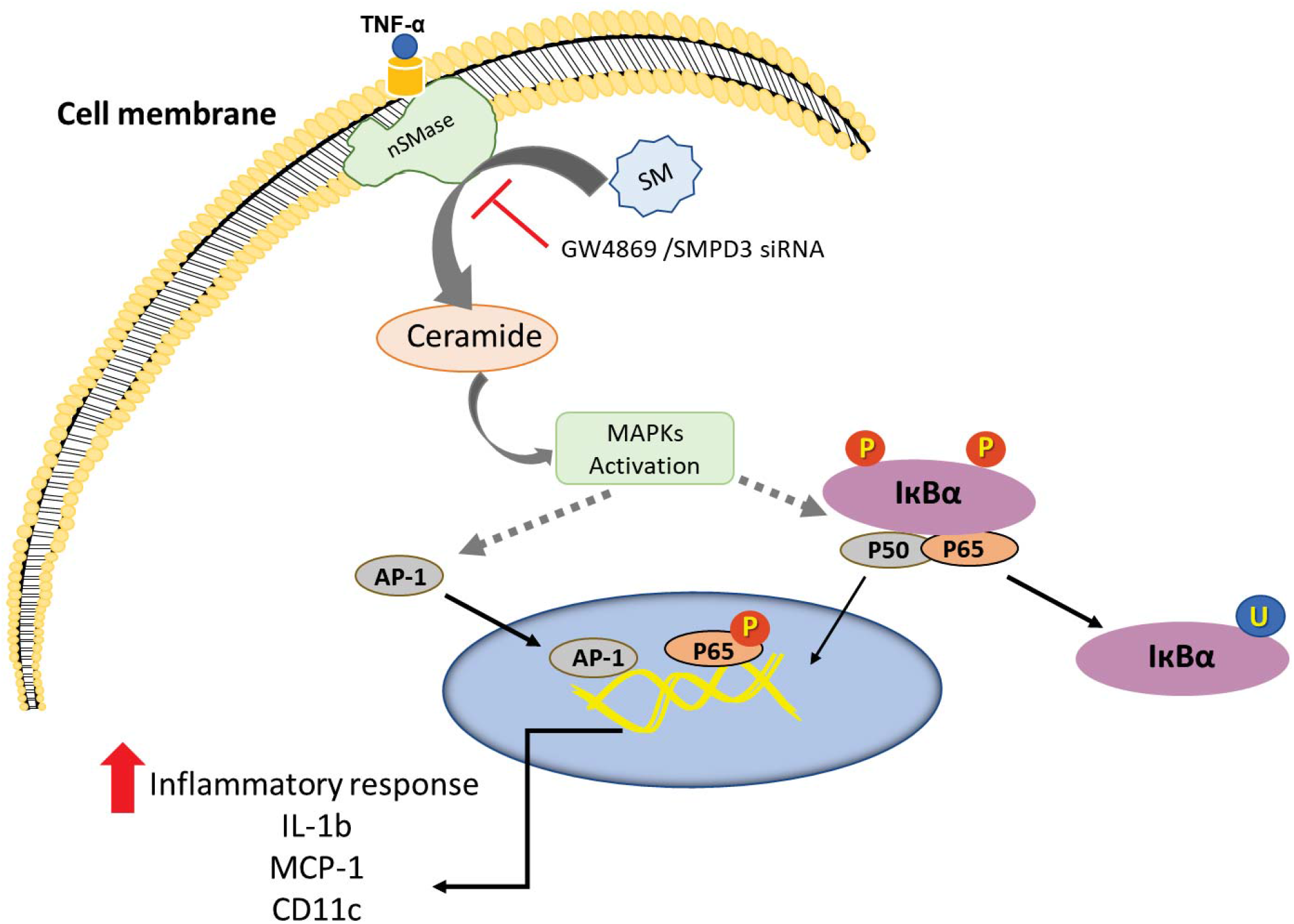
Schematic illustration of the role of nSMase2 in TNF-α mediated inflammatory response in monocytic cells. **T**he pathway highlighted that disruption of nSMase2 activity by GW4869 or SMPD3 suppress TNF-α induced production of inflammatory markers (CD11c, MCP-1, IL-1b) in monocytic cells. SM: sphingomyelin.

## DISCUSSION

Obesity-driven changes of the adipose tissue monocytes/macrophages alters the secretion of proinflammatory cytokines which contributes to the progression of inflammation and the development of insulin resistance (19, 20). Tumor necrosis factor-α (TNF-α), which is overexpressed in obese humans and rodents, plays a central role in metabolic inflammation and insulin resistance. TNF-α in obesity has been considered mainly pro-inflammatory as it activates immune cells, particularly monocytes and macrophages into a pro-inflammatory phenotype, M1 vs M2 (21). However, the etiology of how inflammatory monocytes/macrophages are developed and activated by TNF-α remains unclear. TNF-α activates membrane-associated nSMase2 which cleaves sphingomyelin to produce ceramide, a second messenger activated through the tumor necrosis factor (TNF) receptor (22). In this study we report that nSMase-2 is involved in TNF-α-mediated inflammatory responses of monocytes/macrophages which were characterized by elevated expression of CD11c, an integrin receptor recently shown to regulate adhesion to VCAM-1 on inflamed endothelium. This was accompanied by an increase in the secretion of IL-1b and MCP-1, suggesting monocytic cells activation and having inflammatory phenotype (18, 22, 23). The current results show that inhibiting the activity of the nSMase2 by GW4869 suppressed the TNF-α induced CD11c expression on monocytes. In contrast, the inhibition of aSMase had no effect on the TNF-α mediated expression of CD11c in monocytic cells, demonstrating the participation of a specific sphingomyelinase. Moreover, nSMase2 deficiency prevented TNF-α-induced pro-inflammatory shift in monocytes, suggesting the notion that inflammatory response mediated by TNF-α is suppressed by disruption of nSMase2. Increased expression of CD11c on monocytes occurs in a wide range of inflammatory disorders such as coronary artery disease, asthma, infection, obesity, and T2D (24, 25). It has been reported that high fat diet-induced obesity in mice generates a high numbers of CD11c^+^ monocytes/macrophages and TNF-α (26). Actually, TNF-α-mediated phenotypic shift of monocytes/macrophages in the adipose tissue is a central element of metabolic inflammation and insulin resistance in obese mice.

CD11c^+^ monocytes are actively involved in the production of inflammatory cytokines and chemokines including IL-1b, TNF-α, and MCP-1, pointing to increased migration of the inflammatory monocytes from the circulation into adipose tissue in obesity (27). Our data show that the secretion of the IL-1b and MCP-1 by the TNF-α-activated monocytes was significantly suppressed by blocking of nSMase2. IL-1b and MCP-1 expression in obesity appears to be vital for phenotypic and functional M1 polarization of monocytes/macrophages in response to TNF-α (28). It is evident that monocytes isolated from obese humans with diabetes display an inflammatory phenotype and secrete higher levels of pro-inflammatory mediators such as IL-6, MCP-1, and IL-1β leading in metabolic dysfunction (29). Metabolic dysfunction is implicated in a wide-ranging effect on the immune cells, inflammation and lipid metabolism associated with adipocytes (30). TNF-α activates the plasma membrane nSMase2 that hydrolyzes sphingomyelin to ceramide results its accumulation within the cell (9). Our results show that ceramide level was elevated in TNF-α treated THP1 cells. nSMase regulates inflammatory responses via ceramide production and inhibition or deficiency of nSMase2 reduces inflammation. Lallemand et al. (2018) demonstrated nSMase2 deficiency or inhibition by GW4869 reduces inflammation (31) supporting our findings. Obesity elevates TNF-α expression in adipose tissues (32) and it induces ceramide production via hydrolysis of SM by SMases suggesting that Ceramide is an interlink between overnutrition and the production inflammatory cytokines.

Substantial progress has been made in understanding the ability of TNF-α to activate MAPKs and NF-κB signaling pathways involved in the regulation of several inflammatory cytokines that contribute to the pathogenesis of different inflammatory conditions (33). MAPK signaling molecules interact with each other along with having extensive cross-talk to other inflammatory pathways including NF-kB in orchestration of inflammatory responses (34, 35). It has been reported by our group and others that ERK, JNK and NF-kB contribute to TNF-α-induced IL-8 and CCL4 expression and secretion by fibroblasts (which indicates that MAPK and NF-kB pathways regulate the gene expression of a variety of cytokines, and playing an important role in inflammation. TNF-α-induced CD11c expression, as well as IL-b secretion in the lungs of mice was found to be dependent on p38 MAPK signaling (36). Previous studies have demonstrated that p38 MAPK phosphorylation can result in nSMase2 activation and that it is associated with inflammation stress (37). Consistent with previous studies (18), we found that TNF-α strongly induced phosphorylation of ERK1/2, p38 MAPK, JNK, C-Jun and NF-kB in monocytic cells. Our results showed that blocking of nSMase2 activity inhibits the TNF-α mediated phosphorylation of p38 MAPK, JNK, c-Jun, NF-kB. NF-kB and AP-1 are key transcription factors downstream of TNFR signaling pathways, our results showed that activation of these two transcription factors inhibited by inhibition of nSMase2. These results suggest that nSMase2 may acts upstream of MAPK and NF-kB signaling pathways.

In conclusion, our study unveils a novel effect of nSMase2 on TNF-α mediated inflammatory responses in monocytic cells/macrophages that is dependent on the MAPK and NF-kB signaling pathways. These findings also support a possible association between TNF-α and nSMase2 in obesity that link may contribute to metabolic inflammation.

## Acknowledgements

This work was supported by Kuwait Foundation for the Advancement of Sciences (KFAS) grant RA AM 2016-007, RA 2010-003 to RA and NIH grant GM118128 to YAH.

## Author contributions

FA, ZA, RT, MM, AS performed experiments, analyzed data and participated in writing manuscript. LO, FAM, YAH participated in designing, planning experiments and in critical review and editing manuscript, RA planned, designed experimental work, interpreted data and wrote the manuscript.

